# The Warburg effect as a metabolic niche construction strategy in the tumor microenvironment

**DOI:** 10.64898/2026.06.12.731860

**Authors:** Vaibhav Anand, Herbert Levine

## Abstract

The advantage conferred by the Warburg effect to cancer cells remains unresolved. While extensive research has examined aerobic glycolysis at the intracellular level, emphasizing its possible biosynthetic or ATP-production rate benefits, the fate of the energy-rich lactate exported into the tumor microenvironment (TME) has received comparatively little attention. Yet lactate is far from metabolically inert: it actively shapes the TME, functioning as a toxin to adaptive immune cells and simultaneously promoting growth of several non-cancerous cells in the tumor microenvironment which in turn promote tumor growth. Here, we develop a minimal consumer-resource model incorporating tumor, anti-tumor immune, and pro-tumor stromal populations interacting through shared nutrients, lactate, and metabolic byproducts. We find that lactate secretion imposes a growth-exclusion tradeoff: higher glycolytic allocation suppresses anti-tumor immune cells but simultaneously reduces tumor abundance due to less efficient metabolism. This tradeoff is resolved in the presence of pro-tumor stromal populations capable of metabolic cross-feeding, namely stromal cells that convert lactate into metabolites which feed back to support tumor growth, thereby offsetting the glycolytic cost. The cross-feeding benefit is second-order in population size, requiring a critical tumor mass to operate. Taken together, these results suggest that there may be an ecological advantage of the Warburg effect, contingent on microenvironmental context. Specifically, lactate secretion is beneficial primarily when appropriate stromal partners are present, functioning as a niche-construction strategy that reorganizes the tumor ecosystem in favor of tumor expansion.

## 1 Introduction

Although cancer has historically been viewed primarily as a genetic disease, it is now seen as a systemic and complex disease with several facets, an important aspect of which concerns the immediate surroundings in which the tumor grows, known as the tumor microenvironment (TME). The TME is a complex ecosystem that plays an essential role in disease progression [1–4], and is composed of several cell types. These cells engage in a variety of interactions with each other and the tumor itself through multiple mechanisms, including competition for nutrients, signaling-mediated cooperation, and environmental modification.

Recent research suggests that metabolic interactions constitute a central driver of effective ecological interactions within the TME [5–7]. Understanding the ecological principles shaping this complex ecosystem can help us devise better therapies to tackle cancer. Some of these principles are well established; for instance, tumor and adaptive immune cells compete for key nutrients such as glucose, and such competition can suppress anti-tumor immune activity and thereby promote tumor progression [6, 7]. A crucial metabolic phenomenon that remains incompletely understood even a century after its discovery is the Warburg effect [8–11], in which glucose is converted to lactate even in the presence of sufficient oxygen. Although aerobic glycolysis produces less ATP per glucose molecule than oxidative phosphorylation, its prevalence across many cancer types suggests that it confers important selective advantages [12–15]. While Otto Warburg initially proposed that the effect arises from defective mitochondria [16], this hypothesis is no longer accepted [17]. Instead, the Warburg effect is now regarded as a regulated metabolic strategy that supports biosynthesis and proliferation. Several specific hypotheses have been proposed [18, 19], including faster ATP production rates, enhanced carbon diversion into anabolic pathways, and proteome allocation tradeoffs analogous to overflow metabolism in microbial systems [20].

These existing theories which purport to explain the benefits of Warburg effect largely focus on its intracellular benefits. However, they do not account for why an energetically rich resource like lactate, a product of aerobic glycolysis, is dumped into the tumor microenvironment. Lactate is not simply a metabolic waste but instead can play a crucial role in tumor progression. Specifically, it acts as a key regulator of the TME. Lactate accumulates in the extracellular environment and potently reshapes interactions among TME populations [18, 21, 22]. It suppresses anti-tumor immune populations including effector T cells, while promoting immunosuppressive and pro-tumor phenotypes in other cell types [21, 22]. Lactate has been implicated in macrophage polarization toward pro-tumor states [23], supports the persistence of regulatory immune populations that are more tolerant of lactate-rich conditions [21], and contributes to an immunosuppressive milieu maintained by stromal populations such as cancer-associated fibroblasts [24]. More broadly, lactate-mediated environmental modification has been proposed as a major mechanism by which tumor metabolism reshapes the immune landscape of the TME [21, 22].

It is well-established that stromal cells can actively support tumor growth through metabolic coupling. Stromal populations have been shown to provide tumors with alternative nutrients such as alanine [25], and tumor-derived metabolites such as lactate can be taken up and repurposed by stromal cells in ways that ultimately reinforce tumor growth [26]. These observations suggest that the ecological consequences of lactate secretion extend beyond immune suppression to include the active construction of a cooperative environment for the tumor. This resembles the idea of niche construction in ecology where organisms modify their environments to alter growth and fitness [27] and tumors have been speculated to engage in such a strategy [28, 29]. While ecological theory has provided mechanistic understanding of how resource dynamics generate coexistence, cooperation, and competitive exclusion in microbial communities, analogous resource-based models of tumor ecosystems remain limited. In microbial ecology, consumer-resource frameworks incorporating metabolic cross-feeding have successfully explained the emergence of cooperative community structure [30]. Related mechanistic studies in cancer have also emphasized the importance of tumor metabolism and microenvironmental feedbacks in shaping collective behavior [31]. Extending such approaches to the TME offers a principled way to connect intracellular metabolic reprogramming with population-level ecological outcomes.

Here, we develop a minimal consumer-resource model of the tumor ecosystem in which tumor cells convert glucose into lactate, thereby modifying the microenvironment in ways that inhibit anti-tumor immune populations while promoting pro-tumor stromal activity. Within this framework, lactate is treated not simply as a waste product or as a directly metabolized shared resource, but as an ecological mediator that reorganizes interactions among tumor, immune, and stromal populations. We show that lactate production alone does not generically enhance tumor growth. Instead, tumor proliferation is significantly amplified when lactate-mediated environmental modification enables metabolic cross-feeding with pro-tumor populations. Our results provide a new ecological aspect of the Warburg effect, demonstrating how cellular metabolic strategies can reshape ecosystem dynamics and offering an additional reason why cancer cells might adopt this metabolic strategy.

## 2 Model

Inspired by minimal microbial consumer resource models that incorporate metabolic crosstalk to allow cooperation between species [30], we construct a minimal model of metabolism in the tumor microenvironment. In our model, we consider three populations: a tumor population denoted by *T*, an anti-tumor population representing the adaptive immune system denoted by *A*, and a pro-tumor population encompassing several cell types which promote tumor growth denoted by *P*. Such a partitioning of cell types in the tumor microenvironment has been previously established and studied in several contexts [32–34].

We model the resource dynamics among these three classes of cell types in the TME and partition the resources into three classes, *R*^*c*^, *R*^*x*^, and *R*^*r*^. *R*^*c*^ denotes core metabolic resources that are essential for proliferation and are actively consumed by all cell types in the microenvironment. These include glucose which constitutes the dominant carbon source in the tumor ecosystem. *R*^*x*^ denotes toxic metabolic byproducts produced by aerobic glycolysis in the tumor cells. These include lactate, protons, and reactive oxygen species, which together acidify and chemically remodel the extracellular milieu. Elevated concentrations of these metabolites suppress the proliferation and cytotoxic activity of anti-tumor immune populations, while stromal cell types are tolerant of, and in some cases benefit from, this altered chemical environment [21, 22].

Finally, *R*^*r*^ denotes rare metabolic resources that are produced by pro-tumor stromal populations through the metabolic processing of *R*^*x*^ and that confer a disproportionate growth advantage to tumor cells. These include amino acids such as alanine, glutamate, glycine, and arginine, which stromal and cancer-associated fibroblast populations have been shown to secrete and which tumor cells preferentially channel into biosynthetic pathways supporting nucleotide synthesis, redox balance, and proliferation [25, 26]. The enhanced uptake efficiency parameter *g*^*Tr*^ reflects this biosynthetic premium: rare resources yield more proliferative output per unit consumed than core resources alone. In our model, the symbiosis between tumor and pro-tumor cells is mediated through *R*^*r*^. Together, *R*^*c*^ captures shared competition, *R*^*x*^ captures tumor-driven environmental modification, and *R*^*r*^ captures the cooperative metabolic return that closes the ecological feedback loop.

### 2.1 Minimal consumer resource model for the tumor microenvironment

We assume a general form of the population dynamics, specifically

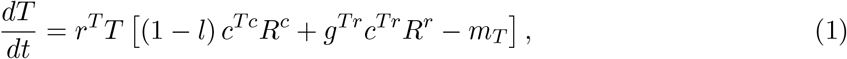

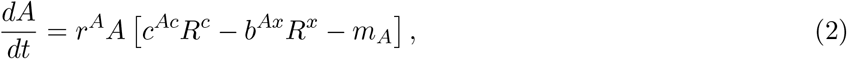

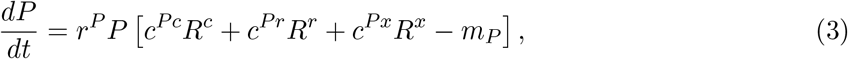

with resource dynamics

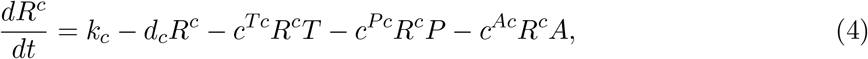

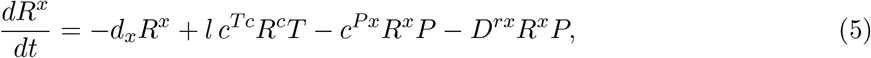

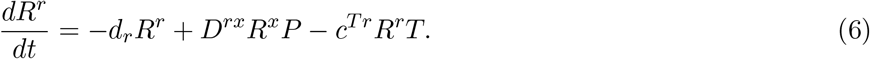

Here *r*^*T,A,P*^ are growth rates, *c*^*ij*^ are resource uptake coefficients for population *i* consuming resource *j*, and *m*_*T,A,P*_ are baseline metabolic costs. The parameter *l* ∈ [0, 1] stands for metabolic reallocation and governs the efficiency of metabolism. Higher *l* indicates a less efficient metabolism and that a greater fraction of consumed resource is redirected into toxic byproduct production rather than growth-supporting metabolism and it is therefore the primary parameter controlling the degree of metabolic reprogramming. The parameter *b*^*Ax*^ captures the inhibitory effect of *R*^*x*^ on anti-tumor cells. The conversion rate *D*^*rx*^ governs the efficiency with which pro-tumor cells transform *R*^*x*^ into *R*^*r*^, and *g*^*Tr*^ *>* 1 reflects the disproportionate growth benefit that rare resources confer on tumor cells relative to core resources alone.

Our dynamical system is completed by the resource equations. *R*^*c*^ is supplied at constant rate *k*_*c*_ and decays at rate *d*_*c*_. Tumor cells convert a fraction *l* of consumed core resources into *R*^*x*^. Pro-tumor cells consume both *R*^*c*^ and *R*^*x*^, the latter at rates *c*^*Px*^ (direct consumption without conversion) and *D*^*rx*^ (conversion into *R*^*r*^). Rare resources are consumed by tumor cells at rate *c*^*Tr*^ and decay at rate *d*_*r*_. We note that while microbial consumer–resource models of cross-feeding incorporate metabolic conversion between resource classes [30], they do not include a toxin produced by one population that simultaneously suppresses competitors and promotes cooperators. This asymmetric ecological role of *R*^*x*^ is a distinctive feature of the present framework.

### 2.2 Reduced two-population model

To isolate the effects of metabolic reprogramming and toxin production, we first study the case where pro-tumor cells are absent. Setting *r*^*P*^ = *c*^*Pc*^ = *c*^*Pr*^ = *c*^*Px*^ = 0 eliminates the pro-tumor population and the cross-feeding pathway, yielding

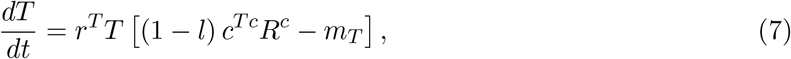

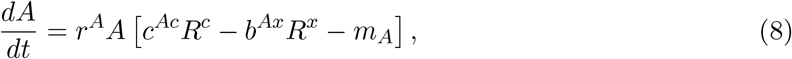

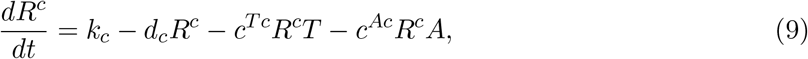

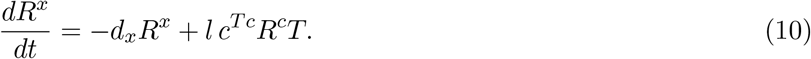

In this reduced system, tumor cells effectively pay a growth cost proportional to *l* while producing a toxin that suppresses anti-tumor cells. This setting allows us to characterize the growth– exclusion tradeoff associated with metabolic reprogramming in isolation, before introducing the cooperative dynamics that arise in the full three-population model.

## 3 Results

### 3.1 Metabolic reprogramming presents a growth-exclusion tradeoff

In the two-species case with only tumor and anti-tumor cells competing for *R*^*c*^, the tumor can reallocate a fraction *l* of its metabolic resources to produce metabolites toxic to anti-tumor cells. The question is whether this strategy benefits the tumor. Simulations in the (*l, b*^*Ax*^) parameter space reveal a clear tradeoff for the tumors (Fig. 2A-D). Higher *l* can exclude the anti-tumor population at lower toxicity *b*^*Ax*^, but the equilibrium abundance of tumor cells decreases with increasing *l* (Fig. 2A). To see this analytically, we assume fast resource dynamics 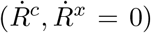 and solve for the equilibrium tumor abundance in the state where it outcompetes the anti-tumor population:

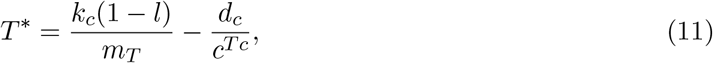

which decreases monotonically with *l*. The stability of this state relies on the issue of whether or not it can be successfully invaded by anti-tumor cells. The invasion exponent of *A* into the tumor-only state *E*_*T*_ can be directly calculated:

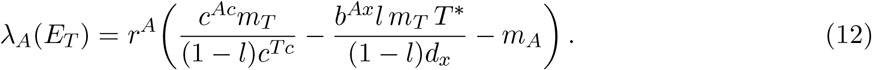

**Figure 1:**
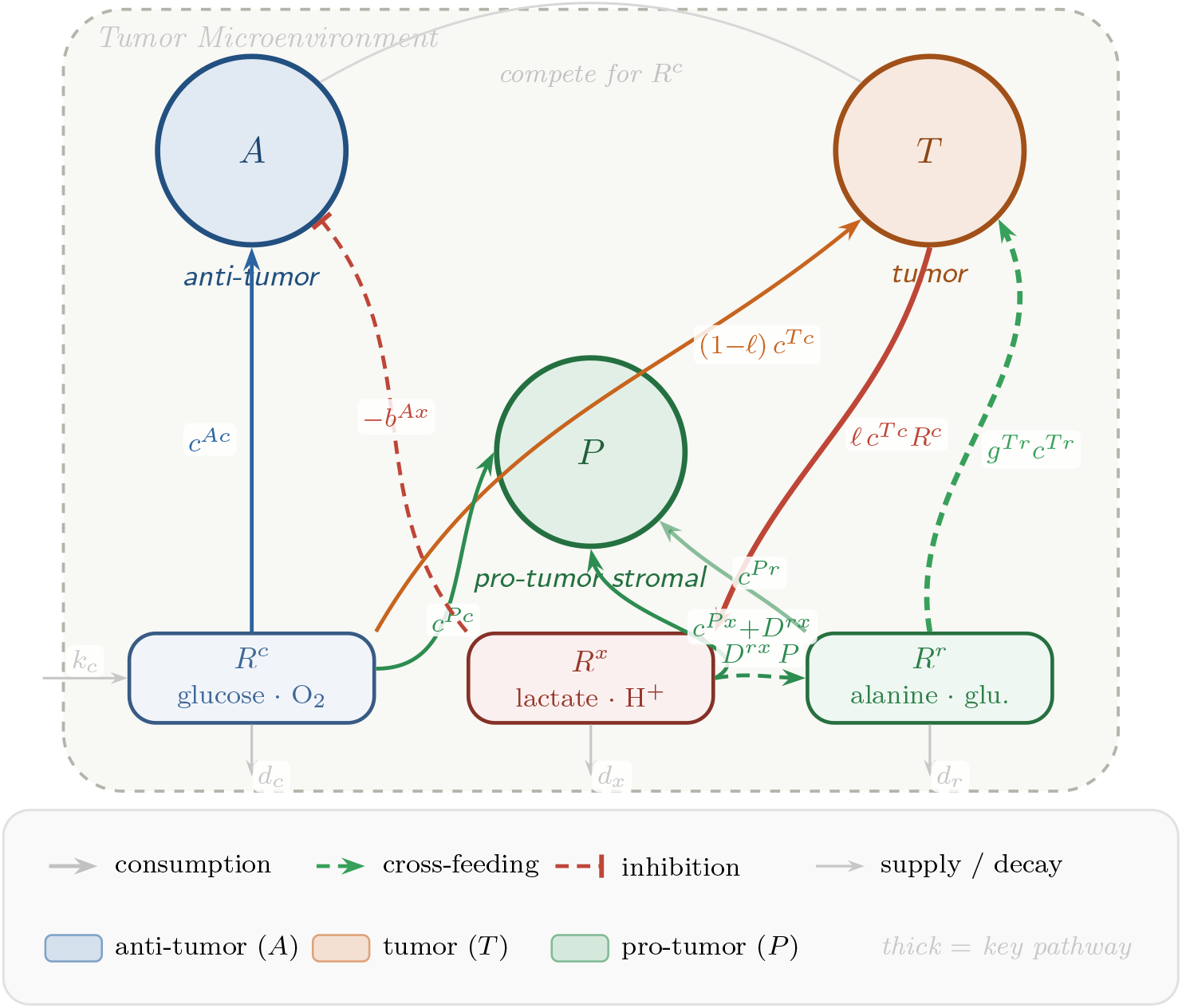
Schematic of the consumer–resource model of the tumor microenvironment. Populations (circles) interact through three resource classes (rounded boxes): core metabolites *R*^*c*^, toxic byproducts *R*^*x*^, and rare cross-fed metabolites *R*^*r*^. Solid arrows denote consumption; dashed arrows denote cross-feeding feedback; the flat-bar indicates inhibition.

**Figure 2:**
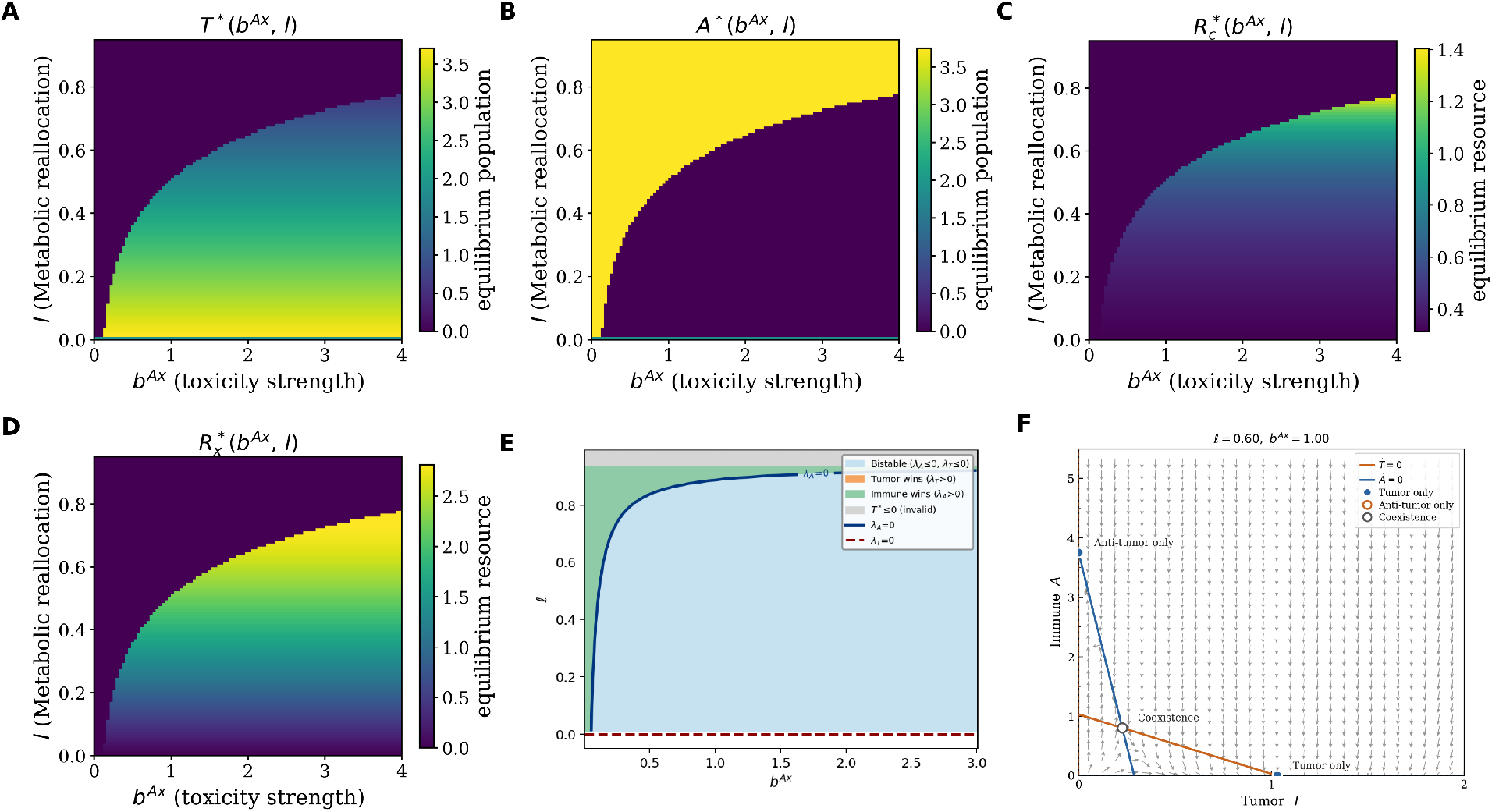
Phase diagrams for the tumor–immune consumer-resource model in the (*b*^*Ax*^, *l*) plane. **(A–D)** Numerically obtained steady-state values of the tumor population *T*\* **(A)**, immune population *A*\* **(B)**, core resource *R*^*c\**^ **(C)**, and lactate concentration *R*^*x\**^ **(D). (E)** Analytic invasion boundary *λ*_*A*_(*E*_*T*_) = 0 *λ*_*T*_ (*E*_*A*_) = 0, demarcating the region where the immune population can invade the tumor-only equilibrium, the region where tumor population can invade immune only equilibrium and the region where neither tumor or immune population can invade each other leading to bistability. The boundaries are derived from quasi-steady-state expressions for *T*\*, *R*^*c\**^, and *R*^*x\**^ evaluated at the tumor-only and immune only fixed points. **(F)** Nullcline and phase portrait of the two species model. Parameters: *r*^*T*^ = *r*^*A*^ = 1, *k*_*c*_ = 1.0, *d*_*c*_ = *d*_*x*_ = 0.2, *c*^*Tc*^ = *c*^*Ac*^ = 0.8, *m*_*T*_ = *m*_*A*_ = 0.25. Initial conditions for figures A-D: *T*_0_ = *A*_0_ = *P*_0_ = 0.05, 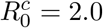, 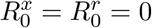.

Setting *λ*_*A*_(*E*_*T*_) = 0 yields the phase boundary shown in Fig. 2E. Higher *l* shifts this boundary, requiring greater toxicity *b*^*Ax*^ to exclude anti-tumor cells.

The other possible state occurs when the only non-vanishing population is that of anti-tumor cells. with this assumption, we find

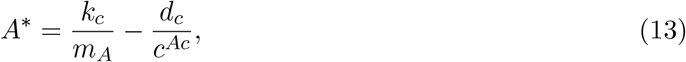

Note that *A*\* is independent of *l* and *b*^*Ax*^, since anti-tumor cells neither produce nor directly interact with *R*^*x*^ in this regime. Now, stability is determined by the inability of tumor cells to invade. The invasion exponent of *T* into the anti-tumor-only state *E*_*A*_ is

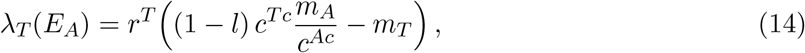

and *∂λ*_*T*_ */∂l <* 0, confirming that higher glycolytic investment strictly reduces the tumor’s ability to invade an immune-controlled environment.

A careful comparison of the two formulas reveals that there exist parameter regions for which both invasions are impossible. This means that the system is bistable, with the actual state depending on the initial conditions (Fig.2E). Finally, it is possible to show that steady-state solutions with both populations being simultaneously present are indeed possible but these are saddle points (Fig. 2F, Fig. S1A,B).

### 3.2 Pro-tumor cells rescue tumor growth through metabolic cross-feeding

We now extend the model to include pro-tumor stromal populations and examine how their presence alters the growth–exclusion tradeoff identified above. We assume throughout that pro-tumor cells grow slower and less aggressively than the tumor and set *r*^*P*^ = 0.5

When pro-tumor cells are present and capable of metabolic crosstalk, the picture changes qualitatively. Pro-tumor cells grow on the toxin *R*^*x*^ that the tumor produces, and in doing so generate *R*^*r*^, which tumor cells metabolize with greater efficiency *g*^*Tr*^. We set *g*^*Tr*^ = 2.5 after exploring the effect of various values of the same (see Fig S1C) This cross-feeding return offsets the growth cost of high *l*, so that tumor abundance is substantially higher across the (*l, b*^*Ax*^) phase space compared to the two-population case (Fig. 3A). In particular, the monotonic decline of *T*\* with *l* seen in the absence of stromal populations is no longer observed: at sufficiently high *l*, the tumor benefits from the enhanced stromal activity it drives, and the cooperative ecological loop more than compensates for the direct metabolic cost of lactate secretion. Apart from the tumor, we also observe an increase in pro-tumor population at higher *l* (Fig. 3C), similar increase is observed in *R*^*x*^, *R*^*r*^ (Fig. 3E,F).

**Figure 3:**
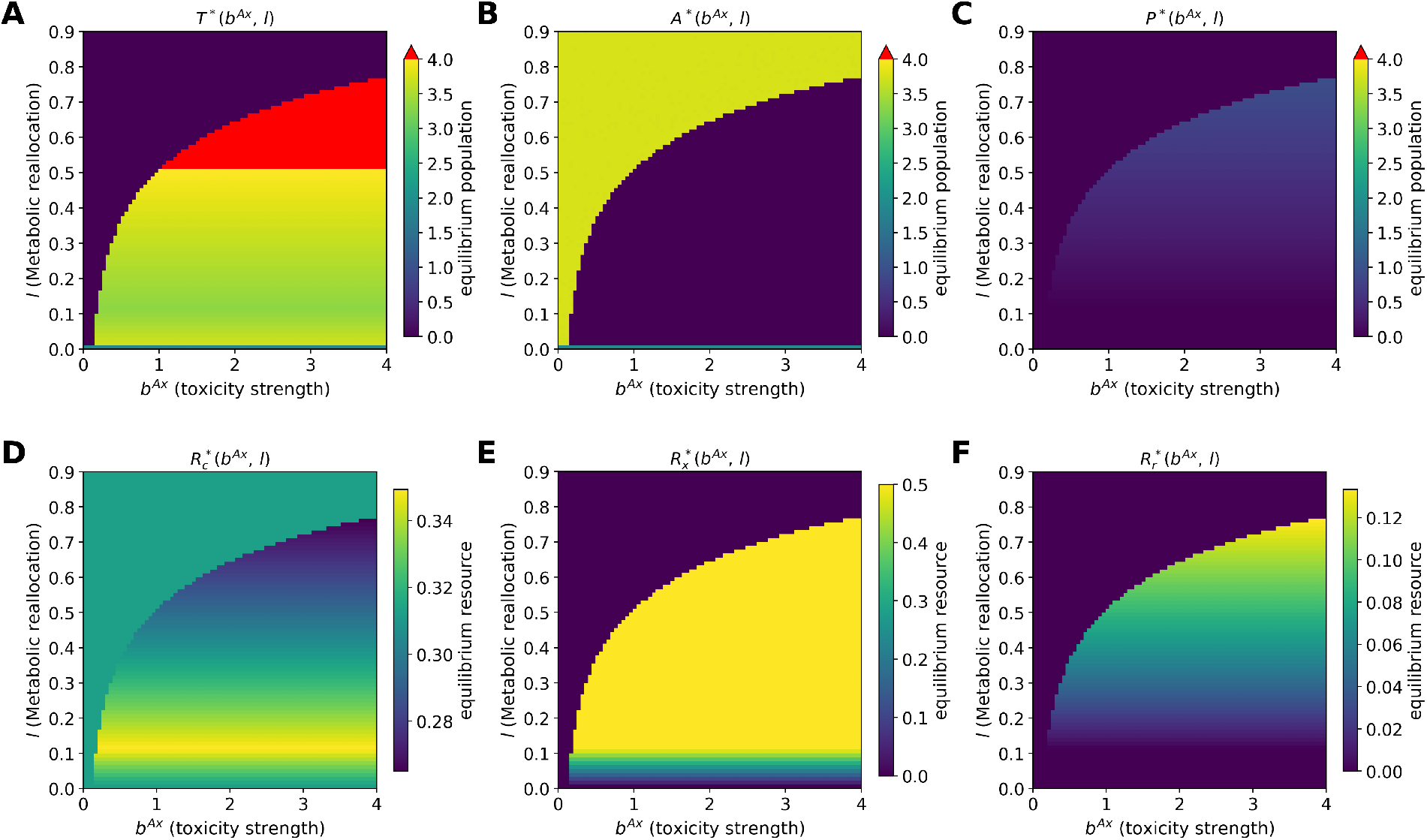
Steady-state phase diagrams for the three-population tumor–immune–stromal consumer-resource model in the (*b*^*Ax*^, *l*) plane. **(A)** Steady-state tumor population *T*\*. **(B)** Steady-state immune population *A*\*. **(C)** Steady-state pro-tumor stromal population *P*\*. **(D)** Steady-state core resource *R*^*c\**^. **(E)** Steady-state lactate concentration *R*^*x\**^. **(F)** Steady-state rare resource *R*^*r\**^. White regions indicate parameter combinations for which a steady state was not reached. Red indicates values exceeding the color scale cap. Parameters: *r*^*T*^ = *r*^*A*^ = 1, *r*^*P*^ = 0.5, *k*_*c*_ = 1.0, *d*_*c*_ = *d*_*x*_ = *d*_*r*_ = 0.2, *c*^*Tc*^ = *c*^*Ac*^ = 0.8, *c*^*Px*^ = 0.5, *m*_*T*_ = *m*_*A*_ = *m*_*P*_ = 0.25, *D*^*rx*^ = 0.8, *g*^*Tr*^ = 2.5. Initial conditions: *T*_0_ = *A*_0_ = *P*_0_ = 0.05, 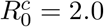, 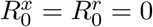.

The Warburg effect, in this setting, is not simply an immune-suppression strategy. It is also the mechanism by which the tumor recruits and sustains its symbiotic partners. Lactate secretion simultaneously weakens anti-tumor populations and selects for pro-tumor cells that return biosynthetically valuable metabolites to the tumor. The net ecological effect of increasing glycolytic allocation is therefore determined by the balance between the direct growth cost, the immune-suppression benefit, and this cooperative metabolic return, a balance that tilts in favor of the tumor only when stromal partners capable of cross-feeding are present.

### 3.3 The cross-feeding benefit is second order and requires a critical tumor mass

The cooperative benefit of cross-feeding does not emerge at arbitrarily small tumor populations. Under the quasi-steady-state approximation for fast resource dynamics, the structure of the feedback is transparent:

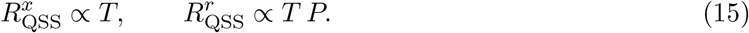

Rare resources arise only through a two-step process: tumor-driven toxin production followed by metabolic conversion by pro-tumor cells. As a result, the cross-feeding return is second order in the populations and vanishes in the limit *T* → 0. Standard invasion analysis, which linearizes around an empty or single-species state, therefore misses this benefit entirely and is not the appropriate tool here.

Instead, we fix an anti-tumor-dominant resident state and introduce varying initial populations of tumor and pro-tumor cells, then track both early growth rates and final abundances. The phase diagram of the final tumor population in the (*T*_0_, *l*) plane (Fig. 4A) reveals that higher glycolytic allocation supports tumor establishment only when the initial tumor population exceeds a critical threshold. Below this threshold, the tumor cannot generate sufficient *R*^*x*^ to sustain pro-tumor growth or initiate the cross-feeding loop, and the anti-tumor population dominates.

**Figure 4:**
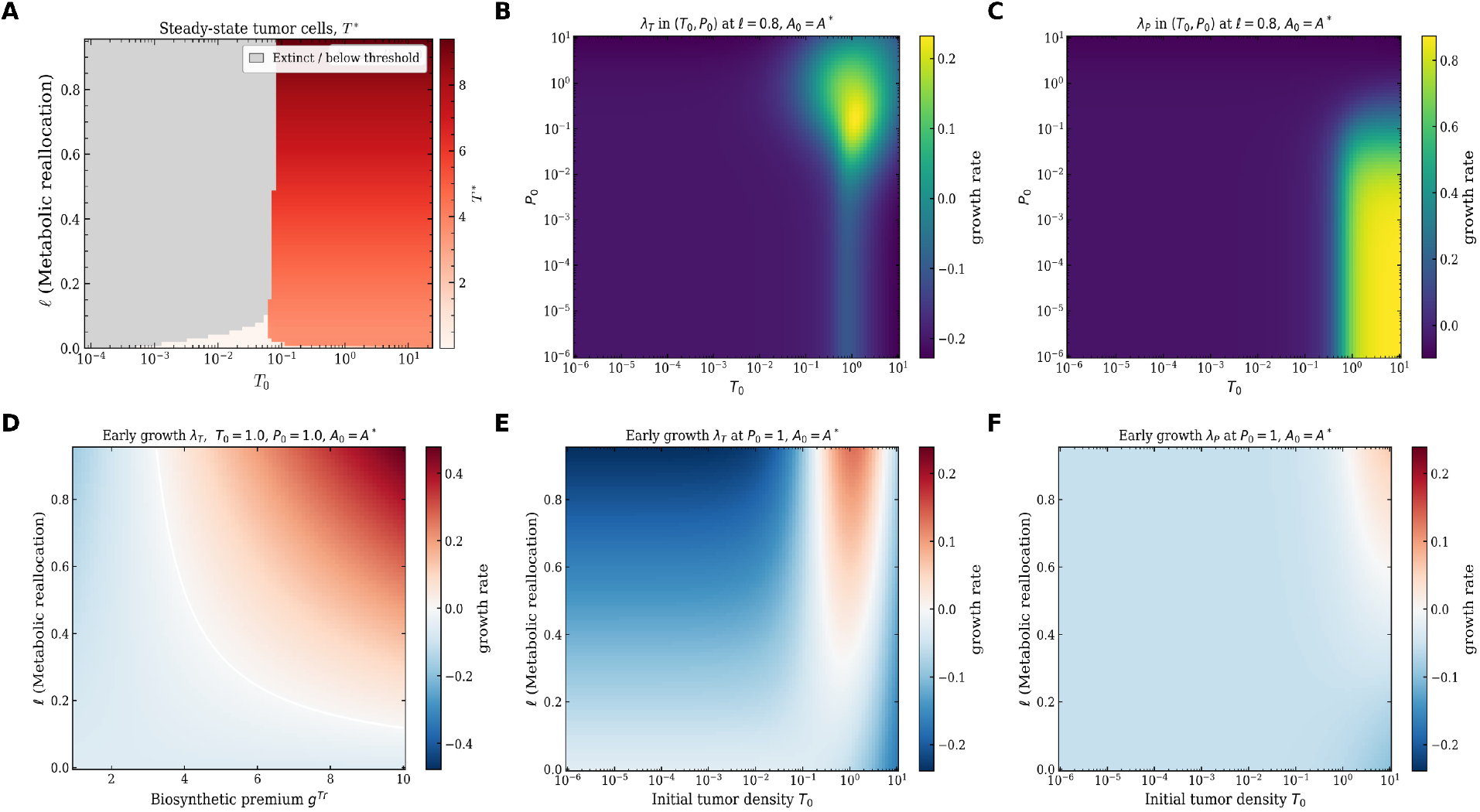
The cross-feeding benefit exhibits an Allee-like threshold. **(A)** Ecological outcomes in the (*T*_0_, *𝓁*) plane at fixed *A*_0_ = 2.0, *P*_0_ = 1.5. **(B, C)** Early-time invasion fitness *λ*_*T*_ and *λ*_*P*_ in the (*T*_0_, *P*_0_) plane at fixed *𝓁* = 0.8, with *A*_0_ = *A*\*. Yellow (blue) indicates positive (negative) per-capita growth. **(D)** Early-time invasion fitness *λ*_*T*_ in (*𝓁, g*^*Tr*^) space. **(E, F)** Early-time invasion fitness *λ*_*T*_ and *λ*_*P*_ in the (*T*_0_, *𝓁*) plane at fixed *P*_0_ = 1, with *A*_0_ = *A*\*. The *T*_0_ axis is logarithmic in all panels. Invasion fitness is estimated as the log-linear slope over *t* ∈ [0, 2]. Shared parameters: *r*^*T*^ = *r*^*A*^ = 1, *r*^*P*^ = 0.5, *m*_*T*_ = *m*_*A*_ = *m*_*P*_ = 0.25, 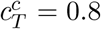, 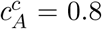, 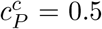, 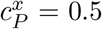, *b*^*Ax*^ = 2, *g*^*Tr*^ = 5, *D*^*rx*^ = 0.8, *d*_*c*_ = *d*_*x*_ = *d*_*r*_ = 0.2, *k*_*c*_ = 1.

We then compute initial per-capita growth rates of both the tumor and pro-tumor populations as a function of their initial abundances. In the (*T*_0_, *P*_0_) plane at fixed *l* = 0.4 (Fig. 4B), tumor growth rates are appreciably positive only when both the initial tumor and pro-tumor populations are sufficiently large, consistent with the cooperative and second-order nature of the cross-feeding return. Pro-tumor cells exhibit analogous behavior (Fig. 4C): growth rates are positive above the critical tumor mass, but turn negative at very high initial pro-tumor abundances, reflecting overshoot beyond the equilibrium capacity.

The dependence of the cross-feeding benefit on the biosynthetic premium *g*^*Tr*^ is illustrated in Fig. 4D, which plots early tumor growth rates in the (*l, g*^*Tr*^) plane. Positive tumor growth rates require a high biosynthetic premium at low glycolytic allocation, but the minimum *g*^*Tr*^ required for net benefit decreases substantially as *l* increases. This confirms that stronger glycolytic investment and a higher biosynthetic premium are complementary requirements for the cross-feeding loop to be ecologically advantageous; the value *g*^*Tr*^ = 5 used in the other panels lies in the regime where the benefit is active at intermediate to high *l*.

Panels E and F of Fig. 4 examine the joint dependence of tumor and pro-tumor growth rates on *T*_0_ and *l* at fixed *P*_0_ = 1. Positive tumor invasion fitness requires both a sufficiently high initial tumor population and *l* ≳ 0.4 (Fig. 4E). Pro-tumor cells display similar but more stringent requirements: positive growth rates demand a higher initial tumor population and a larger glycolytic allocation than the tumor itself (Fig. 4F). This asymmetry reflects the fact that pro-tumor recruitment depends on a sufficient supply of *R*^*x*^, which is itself contingent on the prior establishment of a critical tumor mass. While the results so far examine a tumor-protumor invasion into an established antitumor state, we also observe that the biosynthetic premium *g*^*Tr*^ prevents invasion of antitumor cell types into an established tumor pro-tumor equilibrium (Fig S2A,B).

Taken together, these results indicate that the cross-feeding advantage is contingent on the tumor having already established a sufficient population. The ecological benefit of the Warburg effect through metabolic crosstalk is therefore most relevant in established tumors rather than at the earliest stages.

## 4 Discussion

Metabolic reprogramming (aka the Warburg effect), a phenomenon where cancer cells resort to glycolysis, a metabolic process which is significantly less efficient than the standard oxidative phosphorylation, has long been a mystery with several hypotheses proposed over the years. The ubiquity of this phenomenon led to metabolic reprogramming being classified as one of the key hallmarks of cancer. Current theories include the following: faster response to ATP need [35], the ability to divert resources into biomass production [18, 31] and the now generally discredited notion of mitochondrial damage. A similar phenomenon has been studied in microbial growth as well [20, 36]. All the theories talked about so far have had a cellular focus and an intracellular perspective. A possible role of secretion of energy-rich lactate, a byproduct of glycolosis, remained unexplored.

Over the past decades, cancer progression and tumor dynamics have shifted from a purely intracellular perspective to one which treats the microenvironment as a complex ecosystem and as an organ where cancerous and non-cancerous cells recruited by the tumor interact symbiotically and combat the adaptive immune system through webs of cooperation and competition. This competition and cooperation can actually be coarse-grained as two competing teams/guilds [34]. Therefore, it is natural to consider whether lactate secretion can play an important role in modulating the ecological dynamics.

Motivated by this question, we developed a minimal consumer-resource model that explicitly incorporates glucose uptake, lactate secretion, anti-tumor (immune) populations, and pro-tumor (stromal) populations. Our results reveal a key mechanistic distinction between two roles lactate can play in the tumor microenvironment. Lactate-mediated suppression of anti-tumor cells, by itself, is not sufficient to explain why a more glycolytic strategy should be favored. Increasing allocation toward lactate secretion can inhibit anti-tumor cells, but this does not necessarily translate into higher tumor abundance. When lactate acts only as an inhibitory byproduct, stronger Warburg-like allocation can actually reduce tumor abundance by diverting metabolic flux away from more efficient growth-supporting pathways. Immune suppression alone often will make aerobic glycolysis advantageous.

The picture changes fundamentally once pro-tumor stromal populations are included. Lactate is no longer only a suppressive byproduct; it becomes part of a cooperative ecological loop. Stromal cells benefit from the altered metabolic environment and in turn produce secondary metabolites that support tumor growth. In this framing, the Warburg effect is advantageous not simply because it harms anti-tumor cells, but because it helps construct a tumor-promoting ecological niche. The tumor effectively externalizes part of its metabolic strategy into the surrounding ecosystem and benefits from the response of neighboring cell types.

Our results provide a different way of thinking about the possible functional role of aerobic glycolysis in cancer. Rather than asking only whether glycolysis is the best intracellular strategy for a tumor cell in isolation, one should ask whether it is the best strategy for reorganizing the ecosystem around that cell. In our model, a glycolytic phenotype is not universally advantageous: its benefit appears specifically when the microenvironment contains populations capable of recycling or responding beneficially to its byproducts. The advantage of the Warburg effect is therefore relational rather than purely cell-autonomous.

Much of the theoretical literature on tumor-immune interactions has focused on phenomenological effective interactions and on intracellular metabolic optimization in isolation. The present framework connects these two issues by showing how intracellular allocation decisions propagate outward through shared resources and metabolic byproducts to generate emergent ecological interactions. Consumer-resource models provide a natural bridge between cell metabolism and community-level tumor ecology. We note that the support provided by pro-tumor stromal cells need not correspond to a literal one-to-one conversion of lactate into a tumor-consumable resource. Lactate can function more generally as an environmental signal and ecological modifier that promotes pro-tumor stromal activity, which in turn supports tumor persistence through nutrient provisioning, growth factors, extracellular remodeling, or immune suppression [37–39]. In this sense, lactate acts not only as a metabolite but also as a mediator of ecological feedbacks within the TME. Our model coarse-grains the several distinct ways stromal and myeloid populations may respond to tumor-derived lactate — direct consumption as a carbon source, phenotypic polarization, or secretion of tumor-supporting growth factors [23, 26, 37] into a single metabolically mediated cooperative pathway.

Our model is intentionally minimal, and several additional biological details merit discussion. One important omission is self-toxicity at high lactate concentrations. Experimental and theoretical studies suggest that lactate-rich and acidic conditions can suppress anti-tumor immune activity but can also become harmful to stromal populations and even to tumor cells themselves at sufficiently high concentrations [40, 41]. Our model does not include such self-toxicity explicitly, and instead focuses on the regime in which lactate primarily functions as a tumor-promoting ecological modifier. A natural extension would be to incorporate a nonlinear toxicity cost at high lactate levels, which could generate non-monotonic effects of glycolytic allocation. Spatial structure is a further omission: lactate and glucose gradients are not uniform in real tumors, and prior work has shown that acidification and lactate secretion can generate structured niches with differential localization of tumor and stromal populations [40]. Our framework should therefore be viewed as a coarse-grained ecological model of average interactions rather than a description of local microscopic organization. Thus, there are several directions for future work that follow naturally from these limitations. Incorporating explicit toxicity and saturating effects of lactate would allow both beneficial and harmful roles of acidification to coexist in the same model, potentially generating non-monotonic phase boundaries of the kind seen experimentally. Distinguishing multiple stromal or immune subtypes, especially those known to respond differently to lactate-rich environments would extend the ecological resolution of the framework. A third, and theoretically important, direction would be to move beyond a few interacting populations and develop a framework capable of capturing the interaction heterogeneity present in the tumor microenvironment. Statistical physics based approaches capable of capturing ecosystem-level heterogeneity have been developed in ecology [42, 43]; extending such frameworks to tumor ecology could reveal whether the ecological role of the Warburg effect persists generically in large heterogeneous environments, or whether it depends on particular structural features of the microenvironment.

Taken together, our results argue that the advantage of aerobic glycolysis cannot be understood solely in terms of intracellular efficiency or biosynthetic capacity. Its consequences depend on the ecological structure of the tumor microenvironment. In a purely competitive setting, lactate secretion may offer limited benefit or even reduce tumor abundance. In a structured ecosystem with pro-tumor cross-feeding partners, the same metabolic strategy can become advantageous by reshaping the environment and stabilizing cooperative tumor-promoting states. The Warburg effect, in this view, is not just a metabolic phenotype. It is a form of ecological niche construction.

## 5 Methods

All simulations were performed by numerically integrating the consumer-resource ODEs using the LSODA solver in SciPy, with relative and absolute tolerances of 10^*−*7^ and 10^*−*10^ respectively. Steady states were recorded at *t*_max_ = 4000 and verified by checking that max_*i*_ 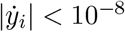 at the final time. Phase diagrams were computed by sweeping two-dimensional parameter grids of 81 × 81 points and recording final-time population and resource abundances at each grid point. Early-time invasion fitness was estimated as the log-linear slope of population trajectories over *t* ∈ [0, 2]. All specific parameter values and initial conditions are reported in the figure captions.

## 6 Acknowledgements

VA and HL acknowledge support by NSF-PHY2019745.

## 7 Author Contributions

VA and HL designed the research, VA did the calculations and both authors wrote the manuscript.

## 9 Appendix

### 9.1 Quasi-steady-state reduction

We consider the limit in which resource dynamics are fast compared to population dynamics, setting

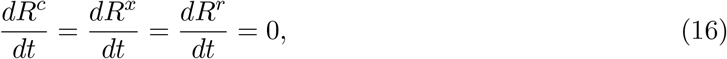

and solving algebraically for the steady-state resource concentrations as functions of the populations (*T, A, P*).

Setting 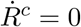 and factoring gives

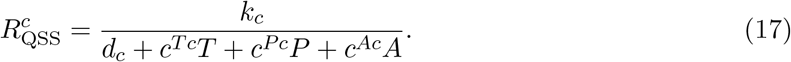

Setting 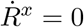 and substituting 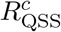 gives

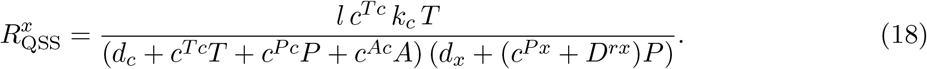

Finally, setting 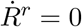 and substituting the expressions above gives

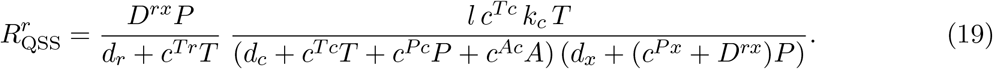

Substituting these into the population equations yields the reduced system

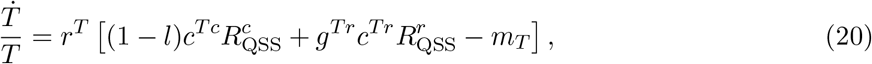

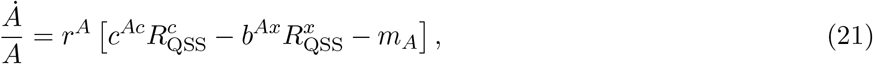

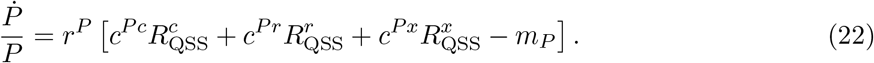

The scaling 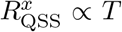 and 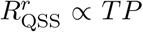 *TP* follows directly, confirming that rare resources arise only through the two-step process discussed in the main text.

### 9.2 Invasion analysis: two-population model

We first carry out invasion analysis on the reduced two-population model with *P* = 0, which isolates the growth-exclusion tradeoff associated with metabolic reprogramming. Let *E* denote a resident equilibrium. The invasion exponent of population *X*_*i*_, *i* ∈ {*T, A*} is

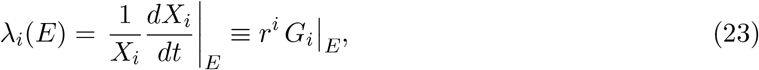

where *G*_*i*_ is the per-capita growth function. A population can invade if *λ*_*i*_(*E*) *>* 0.

#### Tumor-only equilibrium *E*_*T*_ = (*T*\*, 0)

With *P* = *A* = 0, the QSS resources reduce to

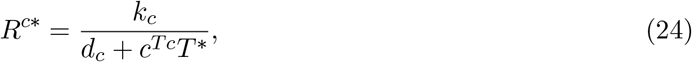

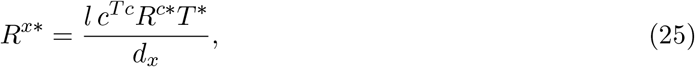

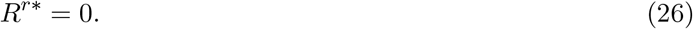

The equilibrium condition *G*_*T*_ = 0 gives *R*^*c\**^ = *m*_*T*_ */*(1 − *l*)*c*^*Tc*^, from which

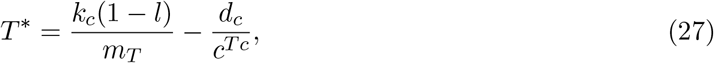

which decreases monotonically with *l*. Substituting into *R*^*x\**^ gives *R*^*x\**^ = *l m*_*T*_ *T*\**/*(1 − *l*)*d*_*x*_. The invasion exponent of *A* into *E*_*T*_ is then

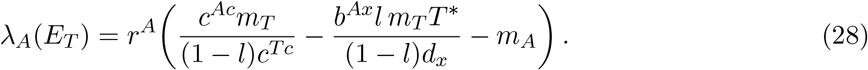

Setting *λ*_*A*_(*E*_*T*_) = 0 gives the phase boundary between tumor-dominant and immune-controlled regimes shown in Fig. 2.

#### Anti-tumor-only equilibrium *E*_*A*_ = (0, *A*^*^)

With *T* = *P* = 0, both *R*^*x*^ and *R*^*r*^ vanish and the equilibrium condition *G*_*A*_ = 0 gives *R*^*c*†^ = *m*_*A*_*/c*^*Ac*^, from which

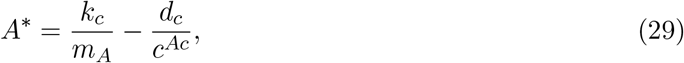

which is independent of *l* and *b*^*Ax*^. The invasion exponent of *T* into *E*_*A*_ is

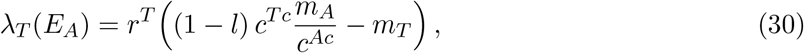

and *∂λ*_*T*_ */∂l <* 0, confirming that higher glycolytic investment strictly reduces the tumor’s ability to invade an immune-controlled environment in the absence of cross-feeding.

### 9.3 Invasion analysis: three-population model

We now consider the full model with *T, A*, and *P*. As discussed in the main text, the crossfeeding benefit scales as 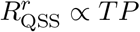 and therefore vanishes in the limit *T* → 0. Standard invasion analysis into an empty or anti-tumor-only state is therefore not informative for the cooperative dynamics. We instead use invasion analysis to characterize which populations can enter a tumor-only ecosystem, and supplement this with simulations at finite initial populations to capture the Allee-like threshold behavior.

#### Invasion of pro-tumor cells into *E*_*T*_

Using the same tumor-only equilibrium *E*_*T*_ = (*T*\*, 0, 0) as above, the invasion exponent of *P* is

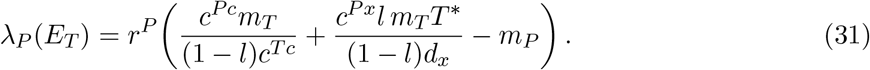

The condition *λ*_*P*_ (*E*_*T*_) *>* 0 determines whether pro-tumor cells can be recruited into a tumor-only ecosystem through toxin availability alone, before any cross-feeding is established. Since *R*^*x\**^ increases with both *l* and *T*\*, higher glycolytic investment and larger tumor populations both facilitate pro-tumor recruitment.

#### A-P subsystem with *T* = 0

When tumor cells are absent, *R*^*x*^ and *R*^*r*^ have no endogenous source and decay to zero at steady state. The dynamics of *A* and *P* reduce to pure competition for *R*^*c*^:

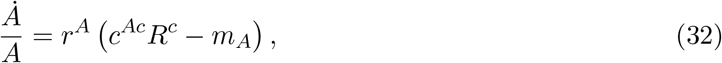

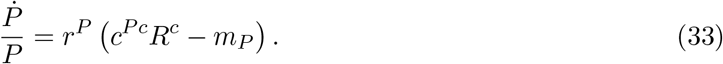

An internal equilibrium with *A*\*, *P*\* *>* 0 requires *m*_*A*_*/c*^*Ac*^ = *m*_*P*_ */c*^*Pc*^, which is non-generic. Generically, one population excludes the other according to which has the lower resource requirement at zero growth. In particular, without tumor-driven toxin production, the cross-feeding and immuno-suppressive pathways are entirely inactive and metabolic interactions play no role in determining community structure. This confirms that the cooperative and suppressive dynamics studied in the main text are contingent on the presence of the tumor.

### 9.4 Jacobian stability analysis of the two-species tumor–immune model

We study the two-population subsystem obtained from the full consumer-resource model by setting *r*^*P*^ = 0 (no pro-tumour stromal cells). The populations are the tumour *T* and anti-tumour immune population *A*, interacting through core resource *R*^*c*^ and toxin *R*^*x*^.

#### 9.4.1 Full two-population equations

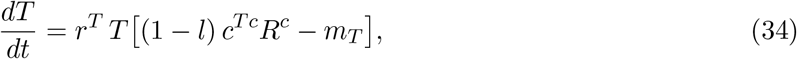

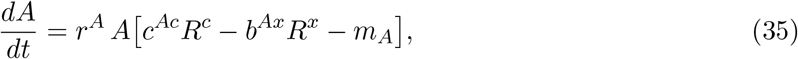

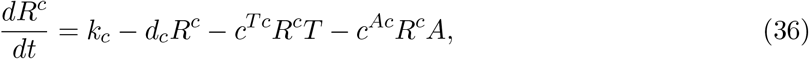

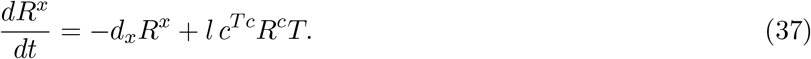

Parameters: *l* ∈ [0, 1] is the glycolytic allocation fraction, *b*^*Ax*^ *>* 0 is the immune-suppression strength of the toxin, *c*^*ij*^ are resource-uptake rates, and *m*_*T,A*_ are metabolic maintenance costs.

#### 9.4.2 Full Jacobian

The Jacobian of the vector field **f** = (*f*_1_, *f*_2_) = (*T G*_*T*_, *A G*_*A*_) is *J*_*ij*_ = *∂f*_*i*_*/∂X*_*j*_. Using the product rule, *J*_*ii*_ = *G*_*i*_ + *X*_*i*_ *∂*_*i*_*G*_*i*_, which simplifies to *X*_*i*_ *∂*_*i*_*G*_*i*_ at any fixed point where *G*_*i*_ = 0. The relevant partial derivatives are

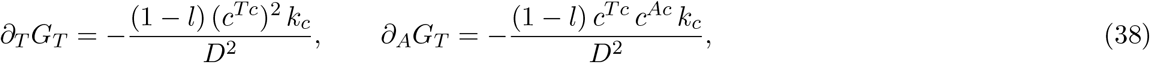

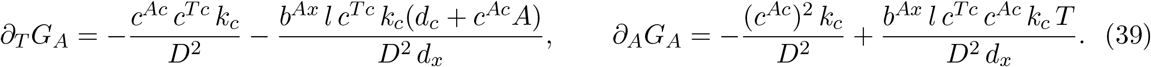

##### Remark 9.1

(Triangularity at boundary equilibria). *At any equilibrium with A* = 0, *the entry J*_10_ = *A ∂*_*T*_ *G*_*A*_ *vanishes. Symmetrically, at T* = 0 *the entry J*_01_ = *T ∂*_*A*_*G*_*T*_ = 0. *Hence J is lower triangular at all three boundary equilibria, and the eigenvalues are simply the diagonal entries*.

#### 9.4.3 Stability of *E*_0_ = (0, 0)

Setting *T* = *A* = 0 gives *D* = *d*_*c*_, so 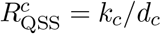 and 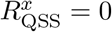. The Jacobian is diagonal:

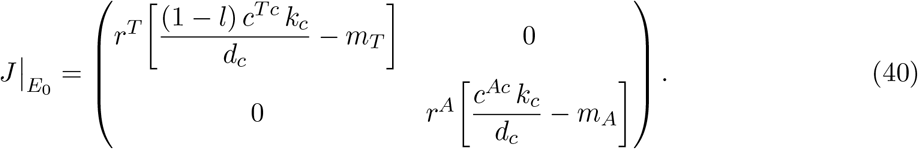

##### Proposition 9.1

*E*_0_ *is unstable whenever both populations can grow from low density, i*.*e*., *when* (1 − *l*)*c*^*Tc*^*k*_*c*_*/d*_*c*_ *> m*_*T*_ *and c*^*Ac*^*k*_*c*_*/d*_*c*_ *> m*_*A*_. *Otherwise it is a saddle*.

The existence of the positive equilibria *E*_*T*_ and *E*_*A*_ already implies *E*_0_ is unstable in the biologically relevant parameter regime.

#### 9.4.4 Tumor-only equilibrium *E*_*T*_ = (*T*\*, 0)

##### Equilibrium coordinates. With *A* = 0, setting *G*_*T*_ = 0 gives

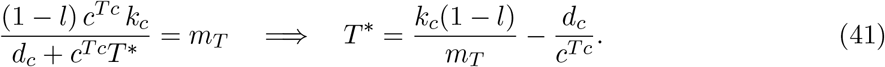

Note that *∂T*\**/∂l* = −*k*_*c*_*/m*_*T*_ *<* 0: the tumour-only equilibrium decreases monotonically with glycolytic allocation, since the Warburg investment directly costs tumour abundance in the absence of stromal cross-feeding. The resource levels at *E*_*T*_ are

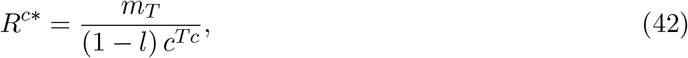

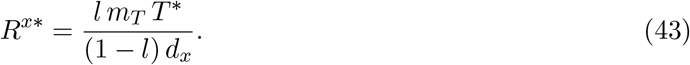

##### Eigenvalues

Since *J* is lower triangular at *A* = 0, the eigenvalues are the two diagonal entries. The first, governing tumour self-regulation, evaluates to

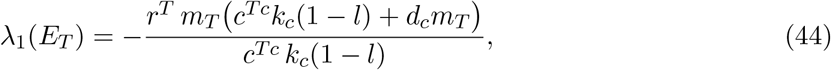

which is always negative when *T*\* *>* 0. The second, the immune invasion exponent, is

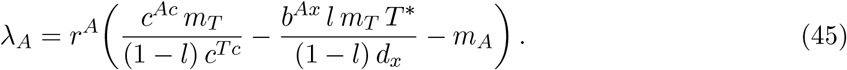

Setting *λ*_*A*_ = 0 gives the phase boundary in the (*l, b*^*Ax*^) plane separating the tumour-dominant from the immune-controlled regime (Fig. 2 of the main text). *E*_*T*_ is a stable node if and only if *λ*_*A*_ *<* 0; it is a saddle when *λ*_*A*_ *>* 0.

#### 9.4.5 Immune-only equilibrium *E*_*A*_ = (0, *A*\*)

##### Equilibrium coordinates

With *T* = 0, both 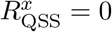 and 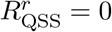. Setting *G*_*A*_ = 0:

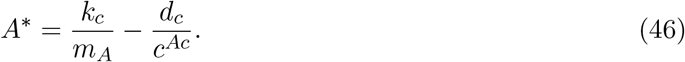

*A*\* depends on neither *l* nor *b*^*Ax*^: when the tumour is absent there is no toxin, and the immune dynamics reduce to a single-species logistic equation in *R*^*c*^ alone.

##### Eigenvalues

The tumour invasion exponent is

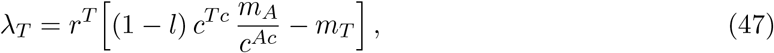

with *∂λ*_*T*_ */∂l* = −*r*^*T*^ *c*^*Tc*^*m*_*A*_*/c*^*Ac*^ *<* 0: higher glycolytic investment strictly reduces the tumour’s ability to invade an immune-controlled environment. The second eigenvalue, governing immune self-regulation, is always negative when *A*\* *>* 0. *E*_*A*_ is stable if and only if *λ*_*T*_ *<* 0.

#### 9.4.6 Bistability

##### Proposition 9.2

(Bistability). *E*_*T*_ *and E*_*A*_ *can be simultaneously stable. This occurs when λ*_*A*_ *<* 0 *and λ*_*T*_ *<* 0, *in which case the outcome depends on the initial condition* (*T*_0_, *A*_0_)

#### 9.4.7 Coexistence fixed point 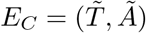

##### Derivation

An interior fixed point satisfies *G*_*T*_ = *G*_*A*_ = 0 with *T, A >* 0. The condition *G*_*T*_ = 0 pins the shared resource independently of *A*:

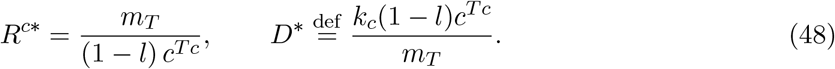

Substituting into *G*_*A*_ = 0 and solving for 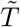 :

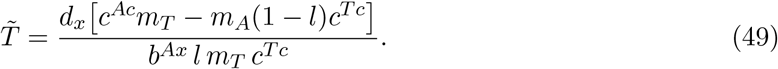

Note that 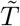 depends on *l* but not on *b*^*Ax*^: in the (*l, b*^*Ax*^) phase plane, varying *b*^*Ax*^ at fixed *l* shifts *Ã* but leaves 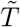 unchanged. Finally, from the denominator constraint *D* = *D*\*:

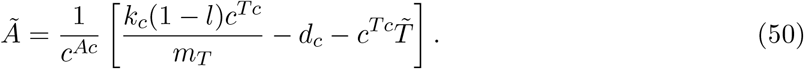

Both 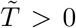 and *Ã >* 0 are required for existence: 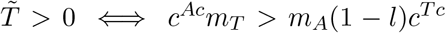 and 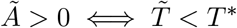.

##### Jacobian at *E*_*C*_

At an interior fixed point *J* is no longer triangular. Using *D* = *D*\* throughout:

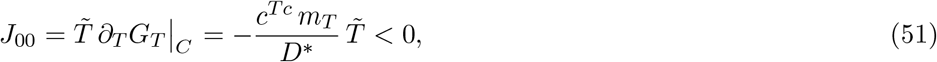

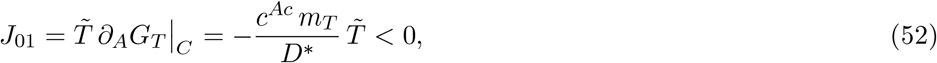

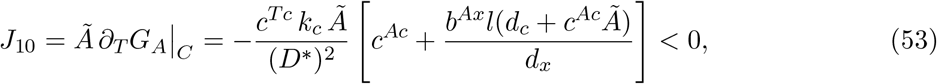

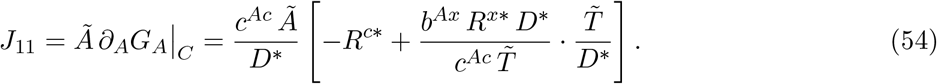

Since *J*_01_ *<* 0 and *J*_10_ *<* 0, their product is positive, so det(*J*_*C*_) = *J*_00_*J*_11_ − *J*_01_*J*_10_ is negative whenever *J*_11_ *>* 0 (strong toxin suppression). The following result holds in general.

###### Proposition 9.3

(Saddle nature of *E*_*C*_). *Suppose E*_*T*_ *and E*_*A*_ *are both stable. If an interior fixed point E*_*C*_ *exists, then* det(*J*_*C*_) *<* 0, *i*.*e*., *E*_*C*_ *is a saddle*.

*Proof*. The Poincaré–Hopf index theorem applied to the positively invariant compact region Ω requires the sum of indices of all fixed points equal the index of any test curve that surrounds those points. Stable nodes or unstable sources have an index +1 and saddles have an index −1. Under the existence conditions for *E*_*T*_ and *E*_*A*_, the trivial equilibrium *E*_0_ has index 1 and any surrounding curve has index +2. Therefore:

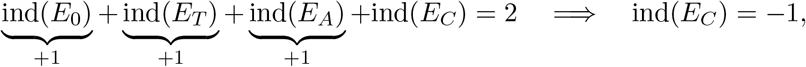

which implies *E*_*C*_ is a saddle and det(*J*_*C*_) *<* 0.

###### Corollary 9.3.1.

*The eigenvalues of J*_*C*_ *satisfy λ*_*−*_ *<* 0 *< λ*_+_. *The stable manifold of E*_*C*_ *(associated with λ*_*−*_*) constitutes the separatrix between the basins of attraction of E*_*T*_ *and E*_*A*_.

### 9.5 Summary of two-species model fixed-point structure

The two-species model is generically bistable in a region of (*l, b*^*Ax*^) parameter space. The Warburg effect (large *l*) shifts the phase boundary to allow immune exclusion at lower toxicity *b*^*Ax*^, but simultaneously reduces *T*\* (eq. 41). Resolution of this growth–exclusion tradeoff requires the three-population model with stromal cross-feeding: the pro-tumour population breaks the bistability by making *E*_*T*_ the global attractor over a much larger region, while the cross-feeding loop offsets the cost encoded in *T*\*.

**Table 1:**
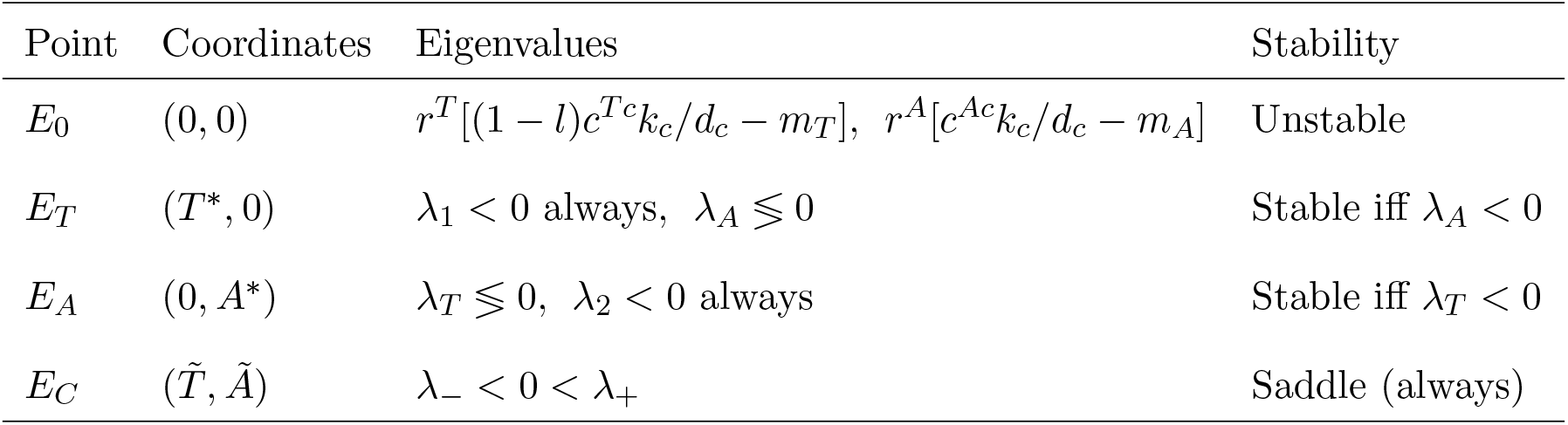
Complete fixed-point catalogue for the two-species QSS-reduced model.

## References

[1] Douglas Hanahan and Robert A Weinberg. Hallmarks of cancer: the next generation. cell, 144(5):646–674, 2011.

[2] Karin E De Visser and Johanna A Joyce. The evolving tumor microenvironment: From cancer initiation to metastatic outgrowth. Cancer cell, 41(3):374–403, 2023.

[3] Lauren MF Merlo, John W Pepper, Brian J Reid, and Carlo C Maley. Cancer as an evolutionary and ecological process. Nature reviews cancer, 6(12):924–935, 2006.

[4] Xueman Chen and Erwei Song. The theory of tumor ecosystem. Cancer Communications, 42 (7):587–608, 2022.

[5] Emily N Arner and Jeffrey C Rathmell. Metabolic programming and immune suppression in the tumor microenvironment. Cancer cell, 41(3):421–433, 2023.

[6] Chih-Hao Chang, Jing Qiu, David O’Sullivan, Michael D Buck, Takuro Noguchi, Jonathan D Curtis, Qiongyu Chen, Mariel Gindin, Matthew M Gubin, Gerritje JW Van Der Windt, et al.Metabolic competition in the tumor microenvironment is a driver of cancer progression. Cell, 162(6):1229–1241, 2015.

[7] Shen Li, Yu Zhang, Huan Tong, Haozhen Sun, Hu Liao, Qingfang Li, and Xuelei Ma. Metabolic regulation of immunity in the tumor microenvironment. Cell Reports, 44(11), 2025.

[8] Otto Warburg, Franz Wind, and Erwin Negelein. The metabolism of tumors in the body. The Journal of general physiology, 8(6):519, 1927.

[9] D Grahame Hardie. 100 years of the warburg effect: a historical perspective. Endocrine-related cancer, 29(12):T1–T13, 2022.

[10] Craig B Thompson, Karen H Vousden, Randall S Johnson, Willem H Koppenol, Helmut Sies, Zhimin Lu, Lydia WS Finley, Christian Frezza, Jiyeon Kim, Zeping Hu, et al. A century of the warburg effect. Nature metabolism, 5(11):1840–1843, 2023.

[11] Ignasi Barba, Laura Carrillo-Bosch, and Joan Seoane. Targeting the warburg effect in cancer: where do we stand? International journal of molecular sciences, 25(6):3142, 2024.

[12] Maria V Liberti and Jason W Locasale. The warburg effect: how does it benefit cancer cells? Trends in biochemical sciences, 41(3):211–218, 2016.

[13] Michelle Potter, Emma Newport, and Karl J Morten. The warburg effect: 80 years on. Biochemical Society Transactions, 44(5):1499–1505, 2016.

[14] Ralph J DeBerardinis and Navdeep S Chandel. We need to talk about the warburg effect. Nature metabolism, 2(2):127–129, 2020.

[15] Rosa Maria Pascale, Diego Francesco Calvisi, Maria Maddalena Simile, Claudio Francesco Feo, and Francesco Feo. The warburg effect 97 years after its discovery. Cancers, 12(10):2819, 2020.

[16] Otto Warburg. On the origin of cancer cells. Science, 123(3191):309–314, 1956.

[17] Deniz Senyilmaz and Aurelio A Teleman. Chicken or the egg: Warburg effect and mitochondrial dysfunction. F1000prime reports, 7:41, 2015.

[18] Matthew G Vander Heiden, Lewis C Cantley, and Craig B Thompson. Understanding the warburg effect: the metabolic requirements of cell proliferation. science, 324(5930):1029–1033, 2009.

[19] B Vibishan, Mohit Kumar Jolly, and Akshit Goyal. Balancing the cellular budget: Lessons in metabolism from microbes to cancer. BioSystems, 256:105571, 2025.

[20] Markus Basan, Sheng Hui, Hiroyuki Okano, Zhongge Zhang, Yang Shen, James R Williamson,and Terence Hwa. Overflow metabolism in escherichia coli results from efficient proteome allocation. Nature, 528(7580):99–104, 2015.

[21] Zhangzuo Li, Qi Wang, Xufeng Huang, Mengting Yang, Shujing Zhou, Zhengrui Li, Zhengzou Fang, Yidan Tang, Qian Chen, Hanjin Hou, et al. Lactate in the tumor microenvironment: A rising star for targeted tumor therapy. Frontiers in nutrition, 10:1113739, 2023.

[22] Joy X Wang, Stephen YC Choi, Xiaojia Niu, Ning Kang, Hui Xue, James Killam, and Yuzhuo Wang. Lactic acid and an acidic tumor microenvironment suppress anticancer immunity. International journal of molecular sciences, 21(21):8363, 2020.

[23] Jordan T Noe, Beatriz E Rendon, Anne E Geller, Lindsey R Conroy, Samantha M Morrissey, Lyndsay EA Young, Ronald C Bruntz, Eun J Kim, Ashley Wise-Mitchell, Mariana Barbosa de Souza Rizzo, et al. Lactate supports a metabolic-epigenetic link in macrophage polarization. Science advances, 7(46):eabi8602, 2021.

[24] Fumimasa Kitamura, Takashi Semba, Noriko Yasuda-Yoshihara, Kosuke Yamada, Akiho Nishimura, Juntaro Yamasaki, Osamu Nagano, Tadahito Yasuda, Atsuko Yonemura, Yilin Tong, et al. Cancer-associated fibroblasts reuse cancer-derived lactate to maintain a fibrotic and immunosuppressive microenvironment in pancreatic cancer. JCI insight, 8(20):e163022, 2023.

[25] Cristov∼ao M Sousa, Douglas E Biancur, Xiaoxu Wang, Christopher J Halbrook, Mara H Sherman, LI Zhang, Daniel Kremer, Rosa F Hwang, Agnes K Witkiewicz, Haoqiang Ying, et al. Pancreatic stellate cells support tumour metabolism through autophagic alanine secretion. Nature, 536(7617):479–483, 2016.

[26] Lifeng Yang, Abhinav Achreja, Tsz-Lun Yeung, Lingegowda S Mangala, Dahai Jiang, Cecil Han, Joelle Baddour, Juan C Marini, Joseph Ni, Ryuichi Nakahara, et al. Targeting stromal glutamine synthetase in tumors disrupts tumor microenvironment-regulated cancer cell growth. Cell metabolism, 24(5):685–700, 2016.

[27] F John Odling-Smee, Kevin N Laland, and Marcus W Feldman. Niche construction. The American Naturalist, 147(4):641–648, 1996.

[28] Irina Kareva. Cancer ecology: Niche construction, keystone species, ecological succession, and ergodic theory. Biological Theory, 10(4):283–288, 2015.

[29] Arig Ibrahim-Hashim, Robert J Gillies, Joel S Brown, and Robert A Gatenby. Coevolution of tumor cells and their microenvironment:”niche construction in cancer”. In Ecology and evolution of cancer, pages 111–117. Elsevier, 2017.

[30] Robert Marsland III, Wenping Cui, and Pankaj Mehta. A minimal model for microbial biodiversity can reproduce experimentally observed ecological patterns. Scientific reports, 10(1): 3308, 2020.

[31] Shubham Tripathi, Jun Hyoung Park, Shivanand Pudakalakatti, Pratip K Bhattacharya, Benny Abraham Kaipparettu, and Herbert Levine. A mechanistic modeling framework reveals the key principles underlying tumor metabolism. PLoS computational biology, 18(2):e1009841, 2022.

[32] Abazar Arabameri, Davud Asemani, and Jamshid Hadjati. A structural methodology for modeling immune-tumor interactions including pro-and anti-tumor factors for clinical applications. Mathematical biosciences, 304:48–61, 2018.

[33] Alexander S Moffett, Youyuan Deng, and Herbert Levine. Modeling the role of immune cell conversion in the tumor-immune microenvironment. Bulletin of Mathematical Biology, 85(10): 93, 2023.

[34] Vaibhav Anand Anand, Mohit Kumar Jolly, and Herbert Levine. Ecological dynamics of pro-tumor and anti-tumorteams in the tumor microenvironment. Physical Biology, 2026.

[35] Matthew A Kukurugya, Saharon Rosset, and Denis V Titov. The warburg effect is the result of faster atp production by glycolysis than respiration. Proceedings of the National Academy of Sciences, 121(46):e2409509121, 2024.

[36] Xin Wang. Overflow metabolism originates from growth optimization and cell heterogeneity. Elife, 13:RP94586, 2025.

[37] Karen G de la Cruz-López, Leonardo Josué Castro-Mun∼oz, Diego O Reyes-Hernández, Alejandro Garćia-Carrancá, and Joaqúin Manzo-Merino. Lactate in the regulation of tumor microenvironment and therapeutic approaches. Frontiers in oncology, 9:1143, 2019.

[38] Ricardo Pérez-Tomás and Isabel Pérez-Guillén. Lactate in the tumor microenvironment: an essential molecule in cancer progression and treatment. Cancers, 12(11):3244, 2020.

[39] Sihan Chen, Yining Xu, Wei Zhuo, and Lu Zhang. The emerging role of lactate in tumor microenvironment and its clinical relevance. Cancer letters, 590:216837, 2024.

[40] Carlos Carmona-Fontaine, Vanni Bucci, Leila Akkari, Maxime Deforet, Johanna A Joyce, and Joao B Xavier. Emergence of spatial structure in the tumor microenvironment due to the warburg effect. Proceedings of the National Academy of Sciences, 110(48):19402–19407, 2013.

[41] Milad Shamsi, Mohsen Saghafian, Morteza Dejam, and Amir Sanati-Nezhad. Mathematical modeling of the function of warburg effect in tumor microenvironment. Scientific reports, 8 (1):8903, 2018.

[42] Guy Bunin. Ecological communities with lotka-volterra dynamics. Physical Review E, 95(4): 042414, 2017.

[43] Madhu Advani, Guy Bunin, and Pankaj Mehta. Statistical physics of community ecology: a cavity solution to macarthur’s consumer resource model. Journal of Statistical Mechanics: Theory and Experiment, 2018(3):033406, 2018.

