## Supplemental figures for "The Warburg effect as a metabolic niche construction strategy in the tumor microenvironment"

### Supplementary Material for:
The Warburg effect as a metabolic niche construction strategy in the tumor microenvironment

Vaibhav Anand, Herbert Levine

June 2026

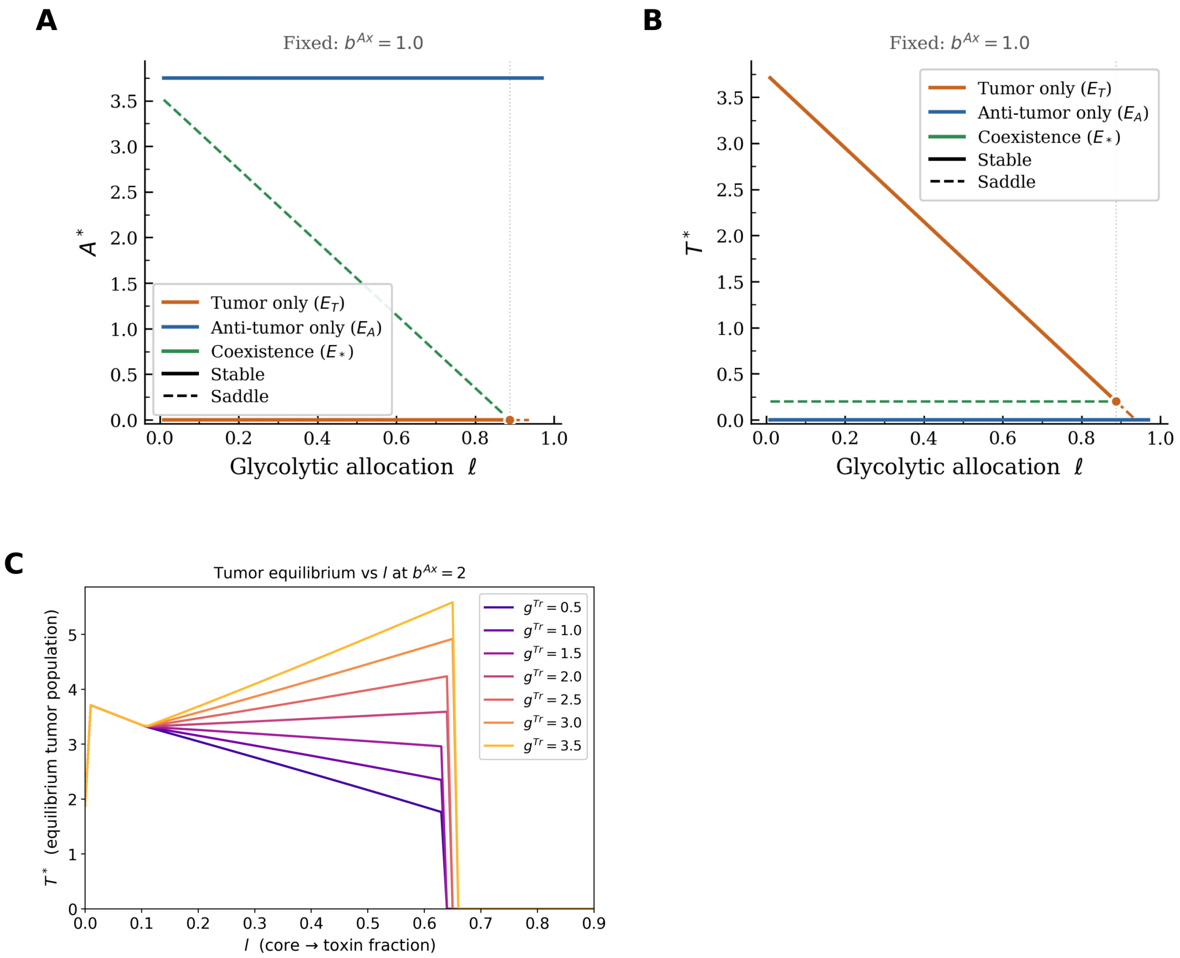


Figure S1: **(A–B)** Bifurcation diagrams showing equilibrium anti-tumor ($A^*$) and tumor ($T^*$) densities as a function of glycolytic allocation $\ell$, with immune suppression fixed at $b^{Ax} = 1.0$. **(C)** Equilibrium tumor density $T^*$ versus $\ell$ for varying rare-resource growth bonus $g^{Tr} \in [0.5, 3.5]$, in the three-species model with stromal cells ($r_P = 0.5$, $D^{rx} = 0.8$). Shared parameters: $r^T = r^A = 1.0$, $m_T = m_A = 0.25$, $c^{Tc} = 0.8$, $c^{Ac} = 0.8$, $k_c = 1.0$, $d_c = d_x = 0.2$.

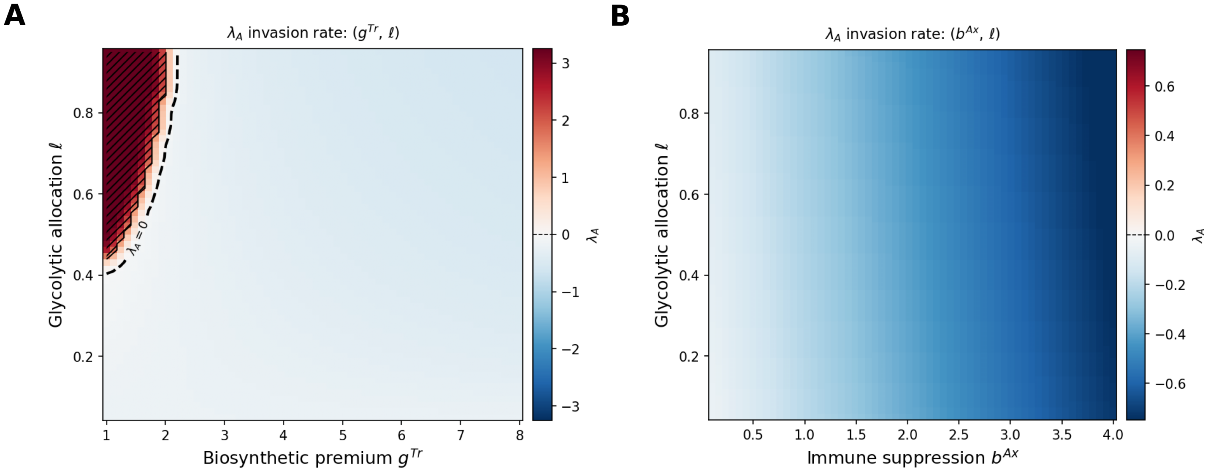


Figure S2: Invasion fitness of anti-tumor cells into the tumor–pro-tumor equilibrium $E_{TP}$. Heatmaps show the invasion exponent $\lambda_A = r_A(c^{Ac}R^c_* - b^{Ax}R^x_* - m_A)$ evaluated at the resident $E_{TP}$ steady state, as a function of **(A)** biosynthetic premium $g^{Tr}$ versus glycolytic allocation $\ell$, and **(B)** immune suppression $b^{Ax}$ versus $\ell$. Red (blue) regions indicate $\lambda_A > 0$ ($\lambda_A < 0$), corresponding to successful (failed) anti-tumor invasion; the dashed black contour marks the $\lambda_A = 0$ invasion boundary. Parameters: $r^T = r^A = 1.0$, $r^P = 0.5$, $m_T = m_A = m_P = 0.25$, $c^{Tc} = 0.8$, $c^{Ac} = 0.7$, $c^{Pc} = 0.5$, $c^{Px} = 0.5$, $c^{Tr} = 0.5$, $c^{Pr} = 0.2$, $k_c = 1.0$, $d_c = d_x = d_r = 0.2$, $D^{rx} = 0.8$; panel-specific sweeps over $b^{Ax} \in [0.1, 4.0]$, $g^{Tr} \in [1.0, 8.0]$, $\ell \in [0.05, 0.95]$, with remaining parameters fixed at base values $b^{Ax} = 1.0$, $\ell = 0.5$, $g^{Tr} = 2.5$.
